# A Systems Biology Analysis of Chronic Lymphocytic Leukemia

**DOI:** 10.1101/2024.03.26.586690

**Authors:** Giulia Pozzati, Jinrui Zhou, Hananel Hazan, Giannoula Lakka Klement, Hava T. Siegelmann, Edward A. Rietman, Jack A. Tuszynski

**Author notes:** These authors contributed equally (Giulia Pozzati, Jinrui Zhou). (GP). (JZ). (HH). (GKL). (HTS). (EAR).

## Abstract

Whole-genome sequencing has revealed that TP53, NOTCH1, ATM, SF3B1, BIRC3, ABL, NXF1, BCR, ZAP70 are often mutated in CLL, but not consistently across all CLL patients. This paper employs a statistical thermodynamics approach in combination with the systems biology of the CLL protein-protein interaction networks to identify the most significant participant proteins in the cancerous transformation. Betti number (a topology of complexity) estimates highlight a protein hierarchy, primarily in the Wnt pathway known for aberrant CLL activation. These individually identified proteins suggest a network-targeted strategy over single-target drug development. The findings advocate for a multi-target inhibition approach, limited to several key proteins to minimize side effects, thereby providing a foundation for designing therapies. This study emphasizes a shift towards a comprehensive, multi-scale analysis to enhance personalized treatment strategies for CLL, which could be experimentally validated using siRNA or small molecule inhibitors. The result is not just the identification of these proteins but their rank-order, offering a potent signal amplification in the context of the 20,000 proteins produced by the human body, thus providing a strategic basis for therapeutic intervention in CLL, underscoring the necessity for a more holistic, cellular, chromosomal, and genome-wide study to develop tailored treatments for CLL patients.

**Author Summary:** Chronic Lymphocytic Leukemia (CLL) is a unique and slowly progressing cancer affecting white blood cells, and research on CLL has highlighted the inconsistency of gene mutations across patients. Using a novel approach that merges statistical thermodynamics and systems biology, this research examines the CLL protein-protein interaction networks to pinpoint proteins integral to the onset of the disease. Betti number (a topology of complexity) estimates, which measure the importance of individual proteins when removed from the network, helped identify numerous potential therapeutic targets, notably within the Wnt signaling pathway, a pathway implicated in various cellular processes and known for its defective expression in CLL. The finding advocates for a multi-target inhibition approach, focusing on several key proteins to minimize side effects, thereby laying a foundation for designing more effective therapies for CLL. This paper emphasizes the potential benefits of a comprehensive study, spanning cellular to genome-wide scales, to design personalized treatments for CLL patients.

## Introduction

Chronic lymphocytic leukemia (CLL) is a type of cancer that affects white blood cells and tends to progress slowly over many years. It is a chronic lymphoproliferative disorder characterized by increased production of morphologically mature but immunologically dysfunctional B lymphocytes. As a result, these cells are unable to fight infections as well as normal white blood cells do [1].

The disease starts developing in the bone marrow, since here leukemia cells survive longer and eventually outnumber normal cells. Then, cells further grow and may spread to other parts of the body including the spleen, the lymph nodes and the liver [1]. Since the growth of leukemia cells is slow, CLL may remain latent for many years before it causes symptoms, and it is usually harder to cure than acute leukemias [1].

From the genetic perspective, CLL is a unique disease with multiple gene signatures. One cohort of patients can exhibit a different gene-signature set than another cohort. Whole-genome sequencing has revealed that TP53, NOTCH1, ATM, SF3B1, BIRC3, ABL, NXF1, BCR, ZAP70 are often mutated in CLL; but, not consistently across all CLL patients [2]–[4]. For example, NOTCH1 is mutated in about 10% of newly diagnosed patients, and in about 15% to 20% of progressive ones. Similarly, SF3B1 is mutated in about 10% of newly diagnosed CLL patients, and about 17% in late-stage disease [2]. Just because a gene is mutated does not mean it will be strongly expressed. One of the goals of our study is to show a molecular thermodynamics approach to determine the most energetically significant pathways supporting a given patient’s CLL initiation and progression. This new molecular systems approach may shed light on optimal treatment for each patient – essentially personalized therapy. Before we present this new methodology, we provide an overview of the known biomarkers for CLL and then a survey of the current treatment options as well as experimental drugs in development.

It is of fundamental importance to obtain information about the patient’s status and prognosis to define therapeutic strategy. There exist several laboratory-based prognostic markers, such as high levels of serum beta2 microglobulin (B2M) and the absolute lymphocyte count (ALC). However, chromosomal aberrations detected using Fluorescent In Situ Hybridization (FISH) serve as the main prognostic tools. The most common aberrations detected in CLL patients are [5]:

- Deletions on the long arm of chromosome 13 (del(13q)): in patients with this aberration the disease progresses slowly.
- Deletions on the long arm of chromosome 11 (del(11q)): this usually occurs among young males and tends to manifest with bulky lymph nodes. It is associated with rapid disease progression and short survival. 11q chromosome contains Ataxia-Telangiectasia mutated gene (ATM) and ATM kinase is responsible for inhibited cell cycle progression in case of DNA damage. Furthermore, ATM kinase acts on p53 by phosphorylating it in order to induce apoptosis. Therefore, when 11q is deleted, this phosphorylation does not occur, and the cell damage cannot be repaired [6].
- Deletion on the short arm of chromosome 17 (del(17p)): results in the loss of TP53 which is the most important prognostic marker in CLL. It is associated with rapid disease progression and resistance to fludarabine chemoimmunotherapy. In addition to the role of TP53 as a prognostic marker in CLL, it is also fundamentally a predictive marker for chemo-immunotherapy, guiding treatment decisions and potentially influencing the response to specific therapies such as fludarabine chemoimmunotherapy [7].
- Immunoglobulin heavy-chain variable region gene (IGHV) mutational status. For prognosis and therapy choice it is important to detect IGHV mutational status since the unmutated state is correlated with low survival.
- Other markers, which are present in a low percentage of newly diagnosed CLL patients, but whose incidence increases in patients who are refractory to fludarabine chemotherapy, are mutations of NOTCH1, SF3B1 and BIRC3. Finally, combining genetics, clinical parameters and biochemistry, the CLL International Prognostic Index (CLL-IPI) is a tool to predict the status of the disease [8].

Wnt signaling is a network of interacting protein pathways which control processes such as cell differentiation, cell cycle regulation, proliferation, apoptosis, cytoskeletal rearrangement, cell polarity, adhesion, motility, migration and invasion and the interaction with the microenvironment [9]. Wnt signaling is correlated with haematopoiesis and is linked with leukaemogenesis of cancers such as CLL [9]. Two Wnt signaling pathways are associated with CLL, namely the Wnt/β-catenin dependent and independent pathways. The Wnt/β-catenin is associated with cell proliferation, homeostasis, cell cycle regulation and thus its malfunction indicated a hallmark of many cancers. Regarding the Wnt/β-catenin independent pathway, the Wnt/PCP (Planar Cell Polarity) is the most important one and it takes place in regulation of cell polarity, migration and invasion. Wnt pathways play a role in CLL pathogenesis and response to treatment. Moreover, the expression of Wnt signaling molecules from Wnt/β-catenin and Wnt/PCP pathways is defective in CLL. For example, ROR1 (receptor tyrosine kinase-like orphan receptor), a Wnt-5 (a Wnt protein) dedicated receptor in the Wnt/PCP pathway, is expressed on the surface of CLL cells and not on the healthy B-cells. Therefore, ROR1 is a sensitive marker of a possible relapse of patients with a more aggressive form of the disease.

The scope of this study extends beyond the traditional single-target silver bullet approach in drug development, acknowledging the intricate network of proteins that drive the pathological transformation of CLL. A systems biology perspective indicates that targeting a manageable group of 5-6 network nodes could be more effective for combination therapy design, considering the potential for serious side effects due to overlapping offtarget interactions. The statistical thermodynamics method applied here aims to identify and hierarchize such targets, which could be inhibited by existing approved or investigational drugs, setting the stage for a more nuanced and personalized treatment approach in CLL.

## Current Treatment Options and Experimental Drug Candidates

Unfortunately, currently available treatments may relieve CLL patients from their symptoms and extend their survival, but still CLL remains incurable [6]. For patients without “active disease” who are asymptomatic or those with early-stage disease, the treatment consists of just a simple observation during which blood counts are performed every three months [6]. For patients with “active disease”, before choosing therapy, the clinical status must be evaluated in terms of general health, characteristics such as TP53 abnormalities or adverse cytogenetics or relapsed disease [6]. Standard treatment has been chemoimmunotherapy with fludarabine, cyclophosphamide and rituximab (FCR). However, it has demonstrated lack of efficacy and it leads to numerous side effects, especially in patients with TP53 or NOTCH1 mutations, unmutated IGHV, deletion of 17p or 11q [8].

Target agents are small molecules that have greater efficacy in patients harboring TP53 mutation or del(17p) whose examples include [6]:

- Bruton tyrosine kinase inhibitors (ibrutinib, acalabrutinib),
- BCL-2 inhibitor (venetoclax),
- Purine analogs (fludarabine, pentostatin),
- Alkylating agents (cyclophosphamide, chlorambucil, bendamustine),
- Monoclonal antibodies (rituximab, ofatumumab, obinutuzumab),
- PI3K inhibitor (idelalisib).

The most common chemotherapy medications used are listed below, together with their main mode of action:

- Fludarabine: a purine analogue and an antineoplastic agent.
- Cyclophosphamide: an alkylating agent.
- Rituximab: a monoclonal antibody which targets the B-lymphocyte antigen CD20 expressed on the surface of B cells.

These three together (FCR) constitute a chemoimmunotherapy treatment:

- Bendamustine: an alkylating agent used along with Rituximab (BR) to form another combination chemoimmunotherapy treatment.
- Chlorambucil: an alkylating agent.
- Ibrutinib is a Bruton tyrosine kinase (BTK) inhibitor. BTK, an enzyme which works for B cell survival and growth, helps delay the progression of cancer. It inhibits CLL cell migration, proliferation and survival [10]. Unfortunately, it presents some side effects such as pneumonia, upper respiratory tract infection, atrial fibrillation, sinusitis, headaches, nausea and many more [10].
- Acalabrutinib is a more selective irreversible BTK inhibitor since it acts just like ibrutinib, but without the side effects involving other kinases [10]. Its most common side effects are headaches, tiredness, low red blood cells, low platelets and low white blood cells [10].
- PI3K (Phosphatidylinositol-3-kinase) inhibitors such as Idelalisib [11], which was FDA approved in 2014 for use in combination with rituximab for treating relapsed CLL [12]. However, Idelalisib is also toxic with nearly 40% of patients having had to interrupt the therapy due to rash or 3-4 grade transaminitis, and pulmonary infections. A PI3Kδ inhibitor called TGR 1202 has better selectivity compared to Idealisib. It was approved for medical use in the USA in February 2021. TGR 1202 reduces the phosphorylation of AKT in lymphoma and leukemia cells.
- Venetoclax binds and inhibits the antiapoptotic protein B-cell lymphoma 2 (BCL-2) [9]. In CLL, inhibition of this pathway has been considered an optimal therapeutic strategy [13]. Use of Venetoclax was approve by FDA in 2016 [13]. Its side effects usually include low levels of white and red blood cells, respiratory infections, diarrhea, nausea, tiredness and tumor lysis syndrome (TLS).
- Sotorasib **(AMG510)** is a highly selective and irreversible inhibitor which binds at an allosteric pocket leading to the trapping of KRAS (Kirsten rat sarcoma virus) in inactive GDP bound state. Note that KRAS transmits signals for growth, division and differentiation to the nucleus of the cell form the outside. KRAS mutations are among the most oncogenic events in carcinomas, including CLL, and the majority of them consist of missense mutation of the 12^th^ codon (glycine). It was approved by FDA in May, 2021. Some of its side effects are diarrhea, nausea and muscles or bone pain [14].
- Adagrasib **(MRTX849)** is an irreversible covalent inhibitor of G12C KRAS mutation that makes a covalent bond to cysteine and binds in the switch-II pocket of KRAS in its inactive GDP state. It demonstrated improved antitumor activity when in combination with vistusertib (an mTOR inhibitor). In clinical trials some patients experienced pneumonitis and heart failure, which led to the interruption of the treatment. Others experienced nausea, fatigue and anemia. This inhibitor is still in clinical trials together with numerous other experimental drugs under development [14].
- Experimental drug candidates also include AKT pathway allosteric inhibitors: ARQ092/miransertib; BAY1125976; MK2206, TAS-117 [15].
- ATP-competitive AKT inhibitors: capivasertib and ipatasertib showed a favorable safety profile along with signs of activity in phase I monotherapy trials [14]. Other AKT inhibitors include the following compounds: Afuresertib **(GSK2110183)**, Uprosertib **(GSK2141795, GSK795**) and Ordidonin **(NSC250682**) [16].

Additionally, various drug candidates are in development with MYC pathway inhibition profiles [17]:

- Compound 361 **(MYCi361, NUCC-0196361**) [18].
- Compound 975 (**MICi975, NUCC-0200975**) [19].
- **MYCMI-6** [20].
- **KSI-3716** and **MYRA-A** [20].
- **KI-MS2-008** [20].
- **L755507** [17].

## Systems Biology Background

The conceptual framework for understanding the thermodynamics and energetics of the molecular biology of human diseases from a network biology perspective has been developed over the past decade. Various studies were undertaken to quantify different signaling and metabolic pathways in various cancer types and other diseases using metrics such as network entropy and the Gibbs free energy applied to each specific case [21]–[26]. Here, we will give only a brief summary of this approach. The transcriptome and other -omic (e.g., proteomic, genomic, etc.) measures can represent the energetic state of a cell. Using the word “energetic”, we mean from a thermodynamics perspective. There is a chemical potential between interacting molecules in a cell, and the chemical potential of all the proteins that interact with each other can be imagined forming a rugged landscape, not dissimilar to Waddington’s epigenetic landscape [27], [28].

The method we propose uses mRNA transcriptome data or RNA-seq data as a surrogate for protein concentration. This assumption is largely valid. Kim et al. [29] and Wihelm et al. [30] have shown an 83% correlation between mass spectrometry-generated proteomic information and transcriptomic information for multiple tissue types. Further, Guo et al. [31] found a Spearman correlation of 0.8 in comparing RNAseq and mRNA transcriptome from TCGA human cancer data [32].

Given a set of transcriptome data, a representative of protein concentration, we overlay that on the human protein-protein interaction network from BioGrid [33]. This means we assign to each protein on the network, the scaled (between 0 and 1), transcriptome value (or RNAseq value). From that we can compute the Gibbs free energy of each protein-protein interaction using the mapping relation:

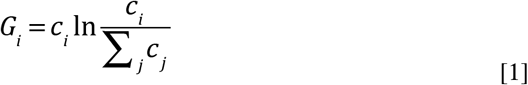

where c_i_ is the “concentration” of the protein i, normalized, or rescaled, to be between 0 and 1. The sum in the denominator is taken over all protein neighbors of i, and including i. Therefore, the denominator can be considered a degree-entropy. Carrying out this mathematical operation essentially transforms the “concentration” value assigned to each protein to a Gibbs free energy. Thus, we replace the scalar value of transcriptome to a scalar function – the Gibbs free energy.

The above equation is derived from a well-known concept in chemical thermodynamics [34]. A biological cell, or a group of cells (a tumor) exist in a complex chemical balance produced by a network of interacting molecular species ranging from small molecules to some very large molecules on the order of hundreds to thousands of Daltons. The molecular concentration-balance in this network is the Gibbs free energy G. This thermodynamic quantity is typically expressed in the context of systems kept at a constant temperature and pressure, where the system can exchange molecules with the environment. For an arbitrary molecular system, the Gibbs function is given as a molar difference [35] in Equation [2]:

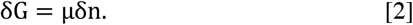

where μ symbolizes the chemical potential, δG is the Gibbs energy and δn is the molar difference (essentially concentration difference). Typically, one writes the chemical potential as:

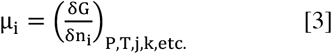

The Equation [3] above assumes that the molar concentrations of other molecular components (other than i) are held constant along with constant temperature and pressure. Using equation [1] and given a network of interacting chemical species, or proteins, and given their concentration, we can compute the Gibbs free energy for a single protein in the PPI.

The Gibbs free energy is a negative number, so associated with each protein on the network is a negative energy well. This results in a rugged energy landscape represented schematically in Figure 1. If we use what is referred to as a topological filtration on this landscape, we essentially move a filtration plane up from the deepest energy well. As the filtration plane is moved up, larger-and-larger energetic subnetworks are captured. For convenience we stop the filtration at energy threshold 32 – meaning 32 nodes in the energetic subnetwork. We call these subnetworks Gibbs-homology networks.

**Figure 1.**
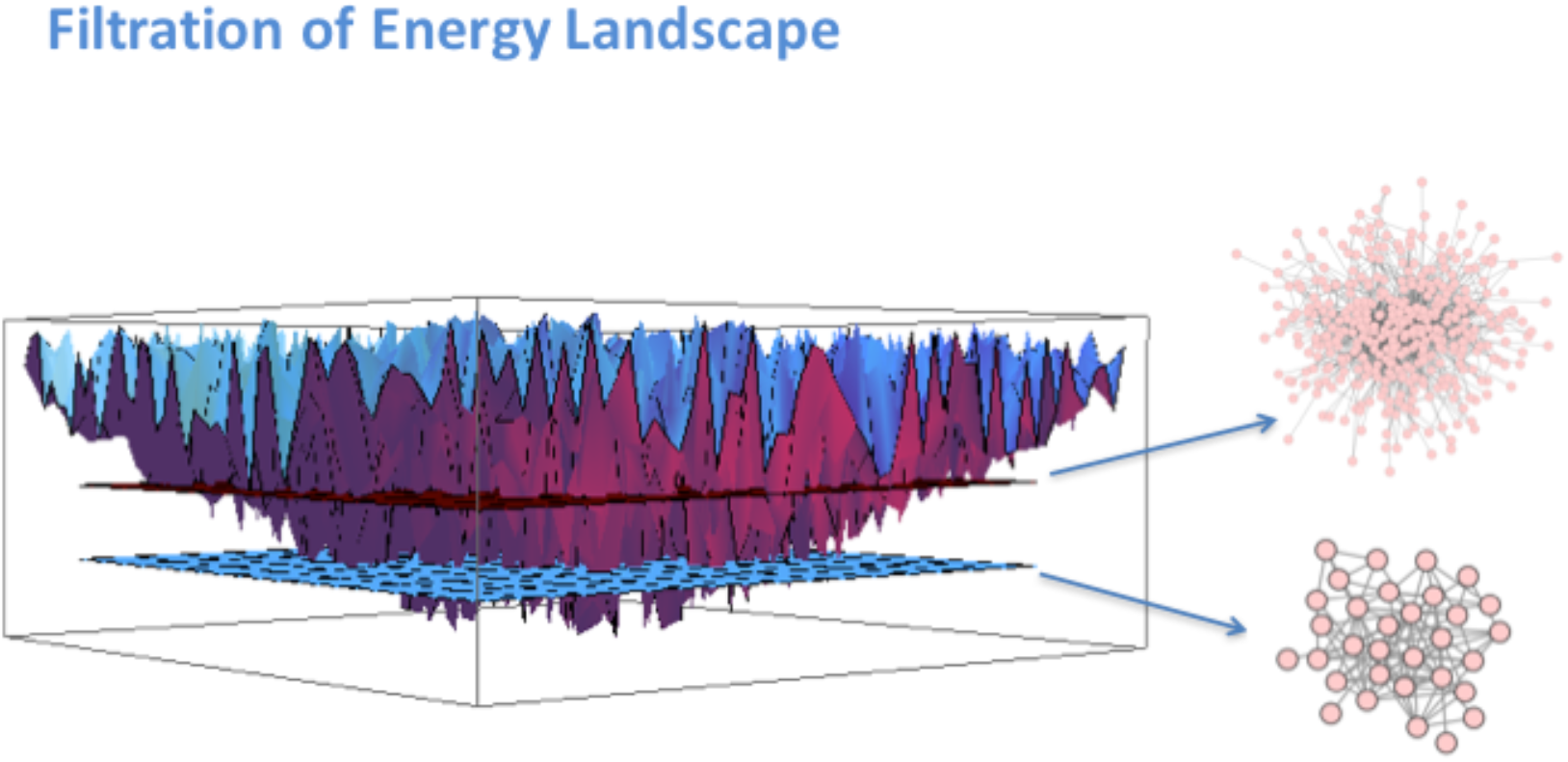
As the “filtration plane” moves up from the bottom, more-and-more nodes are captured in larger-and-larger energetic subnetworks for protein-protein interaction set.

We now compute the Betti centrality, a topological measure, on the 32-node energetic networks as described in Benzekry et al. [23]. The concept is easily described. In networks, there are holes, or rings, of various sizes. In these energetic pathways, protein-protein interaction networks, the proteins form interaction rings. In densely connected, but not fully connected, networks the rings, or holes, may consist of triangles and larger rings of interaction. To find the Betti centrality we ask ourselves: Which protein when removed from the network will change the overall total number of rings the most? The total number of rings is called the Betti number. Given a network G consisting of edges e and vertices v, the Betti centrality is given by Equation [4]:

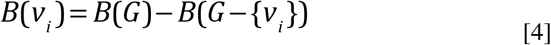

Hence, the difference from the total Betti number B(G) and the Betti number of the network after removing node i, gives the Betti centrality for node i. We compute this for all nodes in the threshold-32 energetic network. Often, there will be two or more proteins in the network that have equivalent Betti centrality.

## Methodology and Datasets

We report on a meta-analysis of 1001 samples from CLL patients and cancer cell lines. This study used online data from GEO [36]: GSE10139, GSE28654, GSE31048, GSE39671, GSE49896, GSE50006, and GSE69034. The data were mRNA expression numbers, all collected using Affymetrix Human Genome Array, HG-U133_2. We also used the human protein-protein interaction network from Biogrid [33]. In particular, we used the dataset downloaded from the BIOGRID-ORGANISM: homo_sapiens-3.5.172.2.

Reiterating the method, we collected the GSE expression datasets, and then the expression value for each gene was overlaid on the human protein-protein interaction network for each protein or node in the network. For each node in the network, we then applied Equation [1] which resulted in the Gibbs energy for that node. This resulted in a rugged landscape similar to Figure 1. Then, the procedure consisted in doing a filtration and computing the Betti number for zero nodes removed and then removing a node and recomputing the Betti number and replacing the node. This removal-computation-replacement procedure resulted in a list of nodes that had the largest impact on complexity of the Gibbs homology network. We finally ranked significant nodes in a Pareto chart for each patient. Pareto charts were prepared at several filtration thresholds, 32, 48, 64, 96.

## Results

Our discussion of the results is presented below, and it follows an analysis of the individual datasets and the research publication associated with it (if present) prior to presenting the meta-analysis Pareto chart and the network graphs.

1. [37] (GSE10137) “A genomic approach to improve prognosis and predict therapeutic response in chronic lymphocytic leukemia”, by Friedman et al. 2009. This was one of the papers with a large table in the Supplementary section. The table consisted of upregulated and downregulated probes indicative of progressive disease; upregulated and downregulated probes indicating chlorambucil resistance; upregulated and downregulated probes indicative of Pentostatin, Cyclophosphamide, and Rituximab signature. An important quote from the paper states that: “Others have previously noted the prognostic significance of cytoskeletal genes and the tumor necrosis factor in CLL. Notably, probes for ZAP-70 did not constitute this genomic signature, although mean expression for ZAP-70 probes in samples from patients with progressive disease was higher than those from patients with stable disease.” The table of genes was parsed from the PDF document and used in our subsequent analysis (discussed below).
2. [3] (GSE28654) “Gene expression profiling identifies ARSD as a new marker of disease progression and sphingolipid metabolism as a potential novel metabolism in chronic lymphocytic leukemia” Trojani, et al. 2012 [3]. A table in the manuscript lists about 65 genes that were selected as being differentially expressed in two cohorts of CLL patients. Of those genes the authors selected 19 genes for PCR analysis because of their significance. Those genes are: ZAP70, ARSD, LPL, ADAM29, AGPAT2, CRY1, MBOAT1, YPEL1, NRIP1, RIMKLB, P2RX1, EGR3, TGFBR3, APP, DCLK2, FGL2, ZNF667, CHPT1, FUT8. An important quote from the paper states that: “In the literature, lists of differentially expressed genes obtained using high-throughput microarray by different laboratories and research centers have often limited overlap [[38], [39]. These differences are matters of important scientific discussions and are imputed, among other causes, to dataset dimensions: small number of subjects (some tens) with respect to the number of variables (tens of thousands of genomic probes in human). … Notably and reassuringly the gene set list (65 genes) emerged from this study showed a substantial (but not quantitated) overlap with results from previously published microarray studies [40]–[43].” The list of 65 genes was incorporated in our subsequent analysis.
3. [40] (GSE31048) “Somatic mutation as a mechanism of Wnt/*β*-Catenin pathway activation in CLL” [40]. In the Supplement to this paper were two large tables listing genes. One table listed from their own study (Wnt pathway) and the other table listed Wnt genes from literature and Websites. Both tables were combined for the study. A quote from the paper: “… our data demonstrate that altered gene expression is indistinguishable between samples with and without mutations.” [41] (GSE39671) “Subnetwork-based analysis of chronic lymphocytic leukemia identifies pathways that associate with disease progression” Chuang et al. 2012 [41]. The Supplemental data only included figures and graphs. No table of gene list. Of note is the quote: “Furthermore, the marker sets identified by different research groups often share few genes in common. Two landmark studies, Rosenwald and colleagues [42] and Klein and colleagues [43] each identify approximately 100 genes that were expressed differentially by CLL cells that use mutated versus unmutated IGHV genes. However, only 4 marker genes were identified in common between these studies.”
4. [44] (GSE49896) “miR-150 influences B-cell receptor signaling in chronic lymphocytic leukemia by regulating expression of GAB1 and FOXP1”, Mraz, et al. 2014 [44]. The following is quoted from their paper: “We identified miR-150 as being the most abundantly expressed miRNA in CLL. However, we observed significant heterogeneity in the expression levels of this miRNA among CLL cells of different patients. Low-level expression of miR-150 associated with unfavorable clinicobiological and prognostic markers, such as expression of ZAP-70 or use of unmutated IGHV (P < .005). Additionally, our data suggest that the levels of methylation of the upstream region of 1000 nt proximal to miR-150 associate with its expression. We demonstrated that GAB1 and FOXP1 genes represent newly defined direct targets of miR-150 in CLL cells. We also showed that high-level expression of GAB1 and FOXP1 associates with relatively high sensitivity of CLL cells to surface immunoglobulin ligation. High levels of GAB1/FOXP1 and low levels of miR-150 associate with a greater responsiveness to BCR ligation in CLL cells and more adverse clinical prognosis.”
5. GSE50006 – no manuscript.
6. GSE69034 – no manuscript.

We created a master list of all genes cited and/or given in the tables associated with the above manuscripts. This list is in the Appendix 2. There were 515 genes total. The list of genes is inputted into the DAVID platform for functional annotation analysis [45], and only 208 genes are found, which indicates that DAVID’s database has annotations for only 208 of those genes. The missing genes might be due to them being less well-characterized, newer discoveries not yet integrated into DAVID’s database, or they might be represented differently in the user’s list compared to DAVID’s nomenclature. To identify genes relevant for a generic condition like “leukemia”, the KEGG [46] and OMIM [47] databases are used to filter and analyze the results such that both are integrated into DAVID. These databases contain curated information about genes related to specific pathways or diseases. By cross-referencing the 208 identified genes with “leukemia” in both KEGG and OMIM, genes whose expression or mutation is linked with the onset, progression, or other aspects of leukemia are pinpointed, aiming in narrowing down potential targets for research, therapeutic development, or further molecular study. Searching that file resulted in the following list AKT1, CTBP1, CTBP2, CTBPA, SMAD4, HDAC1, LEF1, RARA, TCF3, TCF7, TCF7L1, TCF7L2, MYC.

Comparing the PublishedGeneList with our CLLnet96 list, only four were found: MYC, HDAC1, CTNNB1, APP. Two of those, MYC and HDAC1 are known to participate in leukemia. The CLLnet96 list is assembled from all 1001 patients at Gibbs threshold 96. To reiterate the concept of threshold. For any given patient the deepest well in the landscape is, usually the same for all thresholds; but there may be differences based on the expression, and this gives rise to differences in the Gibbs homology network. An energy threshold of, say 32, will result in a network of 32 nodes that are the largest negative energy values. This is called a topological filtration. Using this technique, we can produce one of these 32 threshold networks for each patient. If we do that, and then concatenate the entire list of nodes for each of the patients at this threshold, followed by sorting and discarding redundant nodes in the list, the result will be what we call, CLLnet32 list. By the nature of the filtration, CLLnet32 *χ* CLLnet48 *χ* CLLnet64 *χ* CLLnet96. In words, CLLnet32 is a proper subset of CLLnet48, etc. So, taking the list CLLnet96 will by definition incorporate all others. Comparing our CLLnet96 with the superset of published genes (i.e. PublishedGeneList in the Appendix 2), we find only four that were both lists, MYC, HDAC1, CTNNB1, and APP.

After comparing the PublishedGeneList and the CLLnet96 superset, we then used DAVID, an online bioinformatics resource that allows one to submit a list of genes (or other biological components, e.g. proteins) and it returns important information such as KEGG pathway or OMIM associated with that gene. There are 98 genes in the superlist of CLL96net list. From that analysis we find the following genes to be associated with leukemia (various types): KRAS, GRB2, HDAC1, NPM1, TP53, MYC. While in that superlist, CUL1, TP53, and CTNNB1 is associated with the Wnt signaling pathway.

The published gene lists consisted of two parts. Keeping in mind that although the papers cited above included GSE expression data, most of them did not include tables of genes they identified from their analysis as being important. Instead, they were looking for prognostic markers for disease progression. So, Part 1 of the published gene list consisted of selections identified by the authors from, GSE10137, GSE28654. The combined list consisted of 320 genes. Of those 320 genes, 22 were found in DAVID. Only CEBPA and MYC were found to be associated with any form of leukemia. And CSNK2A1 and MYC were found to be associated with Wnt signaling pathway. When we expand the published list to include GSE321048, which was a focused study on the Wnt pathway and CLL [40], the list expands to 515. Naturally, a huge number of genes were flagged by DAVID as being in the Wnt pathway (78 total). And a smaller subset was found to be associated with some form of leukemia: AKT1, CTBP1, CTBP2, CEBPA, SMAD4, HDAC1, LEF1, RARA, TCF3, TCF7, TCF7L1, TCFL2, MYC. Looking for common genes between the expanded published and our larger list of 96 threshold, we find KRAS, GRB2, HDAC1, NPM1, TP53, MYC, APP, CTNNB1.

At threshold 32, CTNNB1 is best Betti target once out of 1001 patients, but it is present in the threshold 32 networks 326 times. Keep in mind anything found in the 32 threshold is energetically important. So, we find it in 32.5% of the population as a potentially good target for CLL (at 48 threshold 37.9%; at 64 threshold 44.1%; at 96 threshold 58.9%). CTNNB1 is an important gene involved in CLL. It is also an important node in the Wnt pathway [48].

## Results and discussion: Wnt pathway

It is interesting that so many of the authors of the papers cited above did not find overlap among their gene list and other investigators. There was little overlap between those author’s lists until we included the dataset from GSE321048, the Wnt pathway. We speculate that the reason our Gibbs analysis of expression data did not overlap well with other expression data, is that the Gibbs function includes a measure of network entropy (denominator in Equation [1]). Further, many of the genes that are highly expressed, as reported in the literature, are not necessarily mutated based on whole genome sequencing.

Figure 2, shows the Wnt pathway from KEGG. After using the online R-script KEGGraph at Bioconductor it was converted to an edge-list of relevant protein-protein interactions [49].

**Figure 2.**
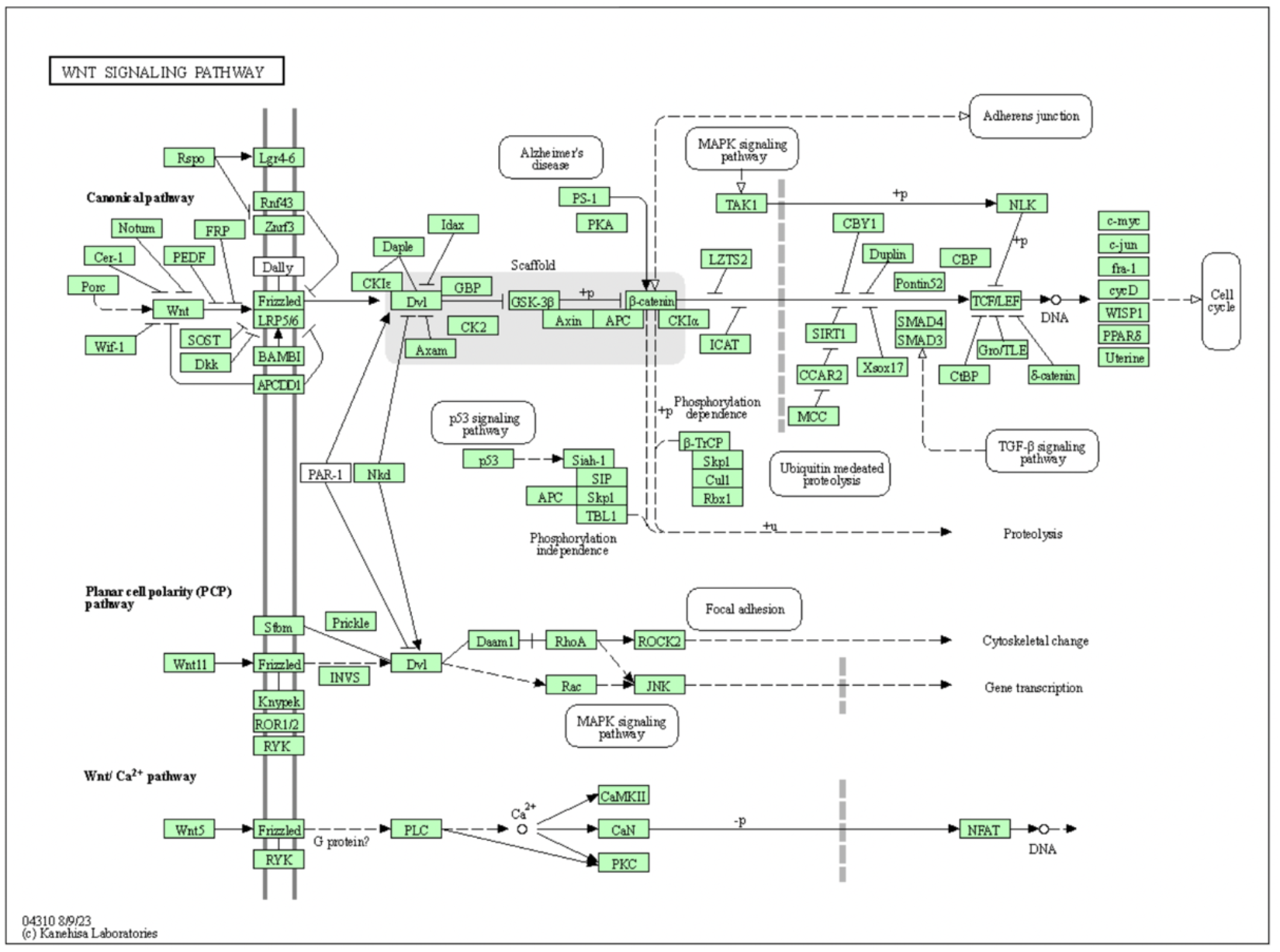
Wnt signaling pathway from KEGG https://www.genome.jp/pathway/hsa04310 [46].

The resulting edge-list was plotted using Cytoscape 3.7 [50]. The PPI network is shown in Figure 3. Two nodes are highlighted. MYC is highlighted and connected to: LEF1, TCF7, TCF7L1 TCF7L2. MYC, as we will see is an important player in Wnt pathway. Also, CTNNB1 has 24 neighboring interactions and has a betweenness of 0.3155, the highest in this network. It also is an important player in the Wnt pathway.

**Figure 3.**
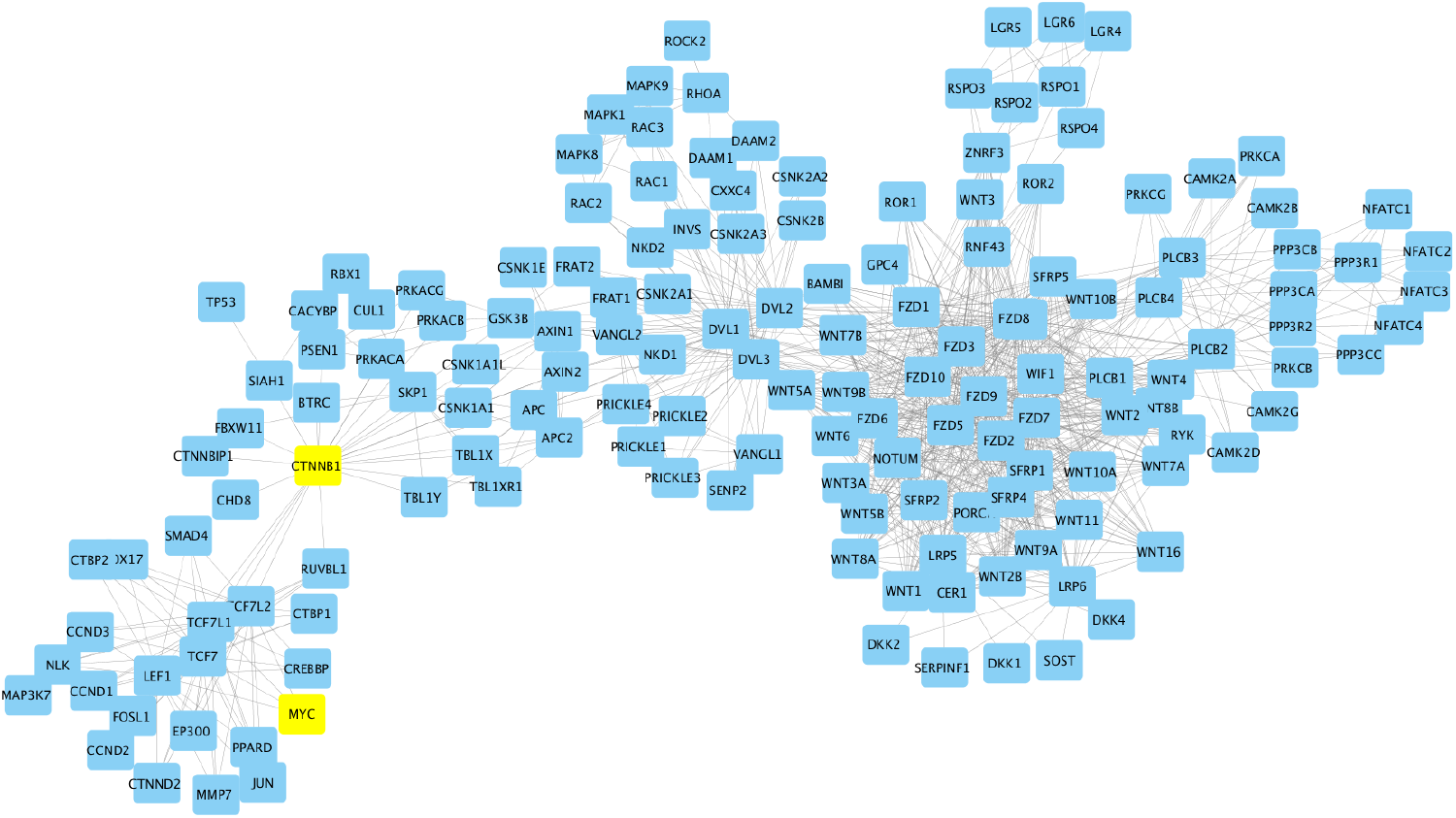
The PPI of Wnt pathway.

As described above, we computed the Betti centrality for the Gibbs homology networks. Figure 4 shows a Pareto chart for the Betti centrality nodes at threshold-48. In our analysis of the 1001 expression samples, CTNNB1 was present as a key Betti centrality node in three samples. Whereas MYC was not present as a key Betti centrality node at threshold-48, but at threshold-32 MYC was present 24 times; 12 times (50%) it was found in dataset GSE30671 which is associated the manuscript by Chuang et al. [41]. This again shows the inconsistency in gene expression values from samples of CLL patients. Of key importance is the fact that RPS15 is a Betti centrality node in 3 patients and RPS15A is a Betti centrality node in 5 patients at threshold-48. This is shown in Figure 4.

**Figure 4.**
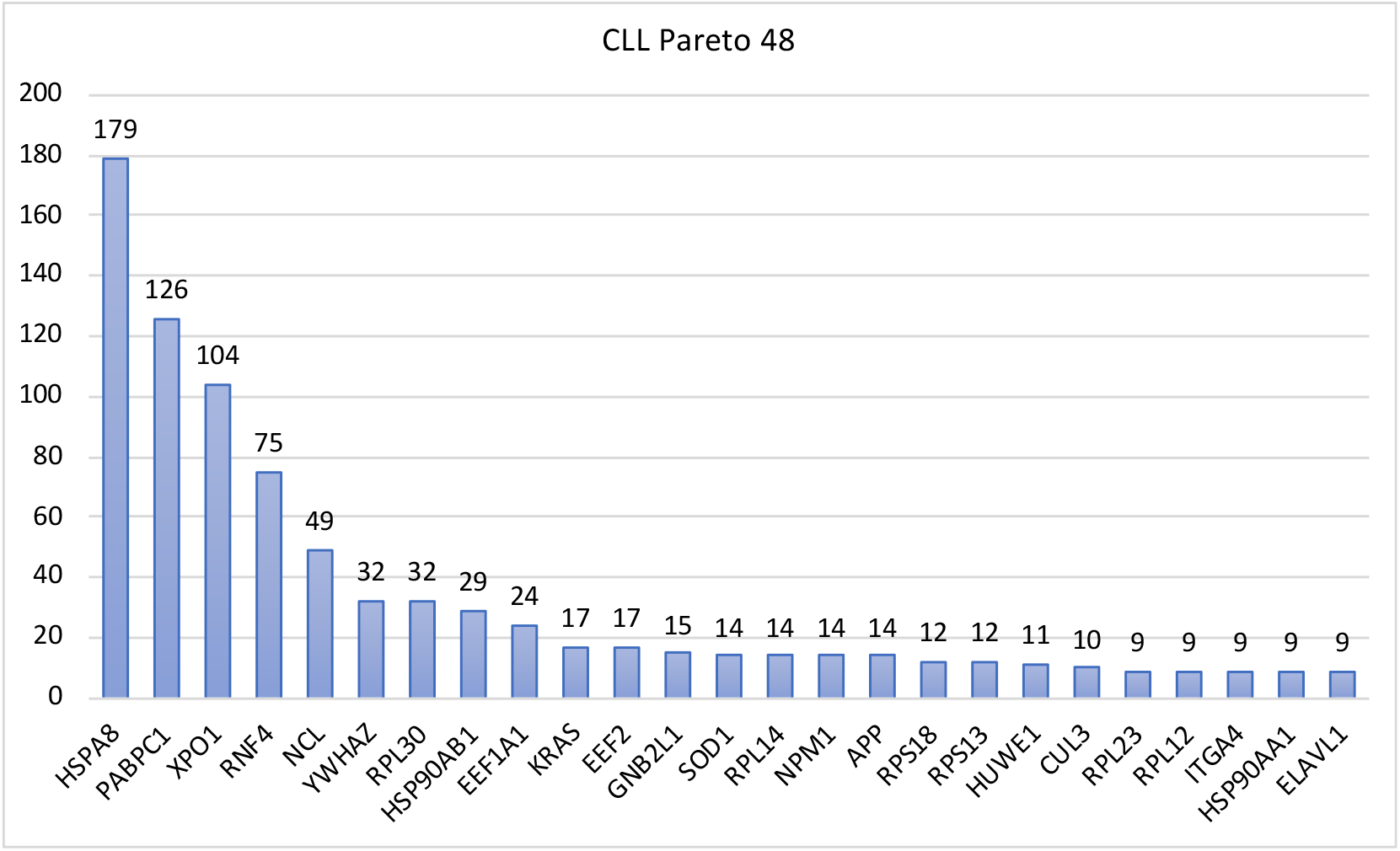
Pareto chart for Betti centrality at Gibbs-homology threshold-48, showing only those with nine or more occurrences.

As shown Figure 3, MYC is an important node in the Wnt pathway. It is directly connected to LEF1, TCF7, TCF7L1, and TCF7L2. Except for LEF1, which is a lymphoid enhancer binding factor, the others are transcription factors. Figure 5 shows a Gibbs homology network at threshold-48 for patient (GSM787065 part of GSE31048 [40]) in which RPS15 is the Betti centrality node. In the network diagram the nodes are in a degree sorted order starting at the bottom with MYC as the highest degree (48) and going around counterclockwise. RPS15 and MYC have been pulled out of the network for easy locating, and MYC with all its first connections have been highlighted in yellow.

**Figure 5.**
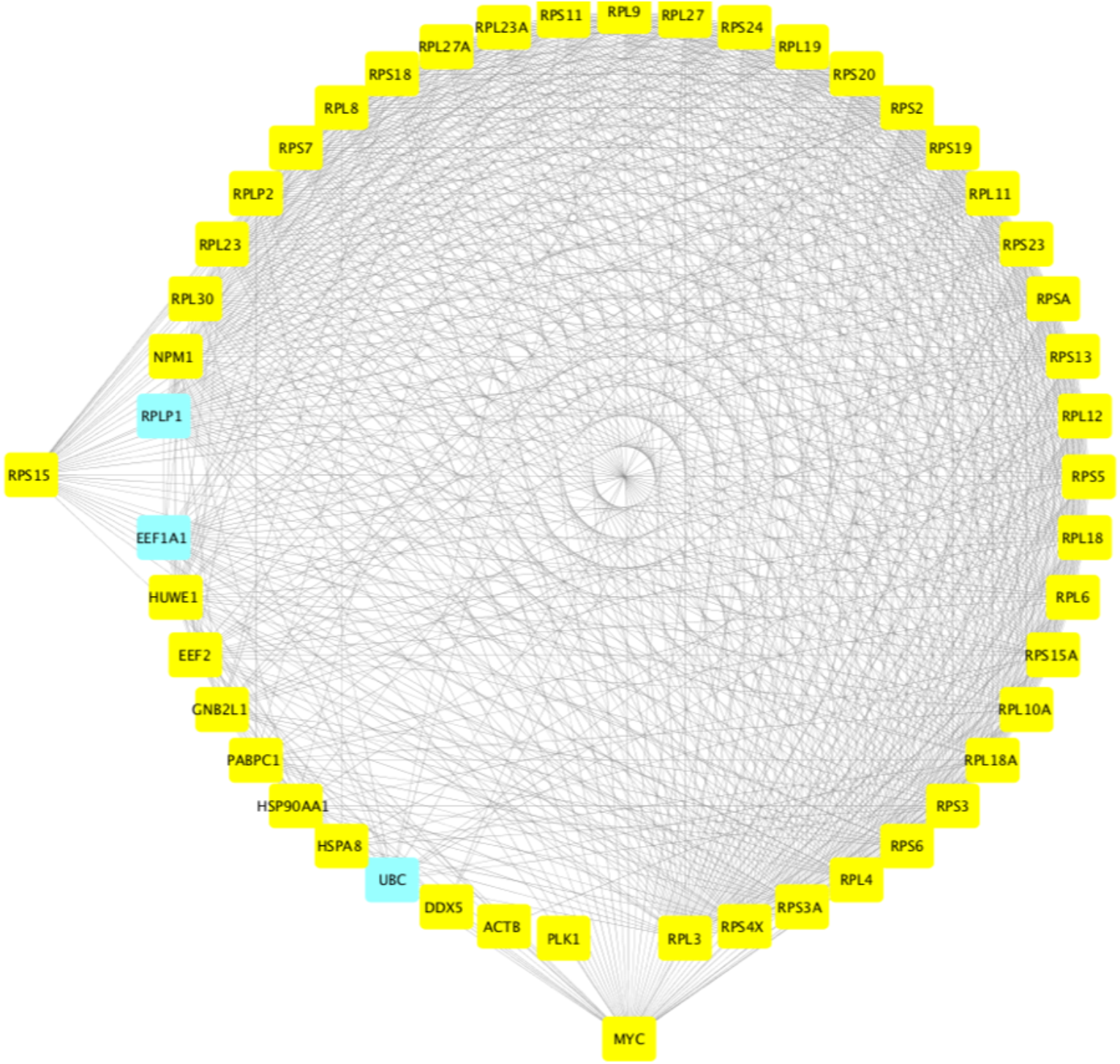
The Gibbs homology network for a patient in which RPS15 has the highest Betti centrality. RPS15 and MYC have been pulled out for easy location. MYC and all its first neighbors are highlighted in yellow.

As we pointed out above, a gene can be mutated, and yet not over-expressed or under-expressed relative to normal. This is likely the main cause for differences in reported transcriptome data from various investigators. What is clear from the literature (e.g. Wang et al.) [40] is that the Wnt pathway is highly important and overexpressed genes in that pathway is often indicative of cancer. MYC is a regulator of ribosome protein synthesis [51] and has been shown to be a key regulator in supporting and maintaining tumorigenesis [52]. For example, Wu, et al. [52] found that inactivation of MYC resulted in some tumors undergoing regression, and mutated RPS15 were identified in almost 20% of CLL patients who relapsed after FCR treatment. These mutations are associated with clinical aggressiveness in CLL along with the mutant RPS15 displaying defective regulation of endogenous p53, which indicates a novel molecular mechanism underlying CLL pathobiology [52]. RPS15 and RPS15A are often overexpressed in CLL [53] and our results confirm this with all (1001 patients) Gibbs-homology subnetworks at threshold-96 showing RPS15 or RPS15A as being an energetically important node.

## Conclusions

We have shown in this study that the genes, BTK, NFkB, JAK/STAT, NOTCH1, BCL2, EEF2, among others play a significant role in the support of CLL. Yet some of them are only rarely studied in the literature because they are not strongly expressed; our research confirms this. Just because a gene or two is mutated does not mean it will be strongly expressed. For example, consider the trisomy of chromosome 7 in colorectal cancers [54]. Using genome-wide chromosome conformation capture (Hi-C), RNA sequencing and protein profiling, along with FISH the authors show that indeed chromosome 7 shows a fair number of upregulated genes relative to healthy cells. However, more strongly they found that chromosome 9 had regions that were very strongly expressed. So, trisomy 7 results in global gene expression changes for colorectal cancer. Might we have something similar going on with CLL patients?

Hi-C analysis is a technique that explores the functional organization of the human nucleome – the folding and unfolding of the chromosomes as the cell goes through its normal processes, including mitosis [55]. Gene topology or arrangement in 3D space affects gene expression, and the 3D topology affects where mutation will occur [56] – [58]. The mathematical technique of processing the Hi-C data involves finding the Laplacian of the “contact matrix” followed by the calculation of the Fiedler vector [59]. The Fiedler vector parses the chromosome into two components: A, the active component, and B, the inactive component. From this it is possible to state, in a high-level way, something about the structure of the chromosome and its function [60]. Since copy number variation closely follows the expression, the function can be deduced from RNA sequencing or transcriptome mapping [61]–[62].

As pointed out above, gene expression found inconsistent sets of genes that were highly expressed and that had high Gibbs-energy. We find the same inconsistent results when we were looking at Hi-C results for CLL patients. Speedy et al. [64] found that BCL2 was strongly implicated in the disease. They also found a disruption at the NFkB-binding site; but, other genes, such as, JAK/STAT, BTK, EEF2 were not mentioned in their manuscript. Beekman et al. [65] found only NOTCH1; Puiggros, et al. [66] found only NOTCH1 and SF3B1 as candidates for high risk of mutation; and Kiefer et al. [67] found NOTCH1 for trisomy 12.

Though the actual causal agent of CLL is not well known, we can speculate that if there is some molecular agent (e.g. herbicide) or an energetic EM signal (e.g. X-ray) it will typically impact the cell only during a specific phase of cell cycle [68]. There are regions of the genome that are more sensitive to alterations due to some specific energy level in the overall molecular network, we call a cell. These mutations are driven by the relevant chemical potential, stereochemistry and Gibbs free energy. We argue that the locations of the relevant genes in the chromosome and the 4D dynamics of the nucleome may suggest a more holistic molecular and cellular approach to understanding CLL and therefore new therapeutic strategies [69]. Building on this notion, the insights from the 4D Nucleome Network [69] elucidate the intricacies of genome organization in space and time. The project underlines the critical role of the genome’s three-dimensional organization in gene regulation. In the context of CLL, the spatial dynamics of chromatin can have a profound impact on gene expression patterns, emphasizing the importance of the genome’s spatial and temporal dynamics in understanding and potentially treating the disease [70]. Elucidating this idea further, the work conducted by Sawh et al. in 2022 reveals that the eukaryotic genome is a multilayered entity, exhibiting intricate organization levels that range from nucleosomes to larger chromosomal scales [71]. These layers undergo significant remodeling across different tissues and developmental stages in C. elegans. It’s noteworthy that advancements in C. elegans research, both imaging-based and sequencing-based, have unveiled the influence of histone modifications, regulatory elements, and broader chromosome configurations in this 4D organization. Specific revelations, such as the physiological implications of topologically associating domains and compartment variability during initial developmental phases, underscore the depth of genome dynamics. These insights provide compelling evidence that understanding such 4D genome organization nuances is crucial for decoding complex diseases like CLL. Interestingly enough, NONE of the genes described in Appendix 1 are in chromosome 13 which often has deletions in about 50% of CLL patients [72]. In Appendix 1 we support our argument for a larger view that CLL genes are widely spread throughout the whole genome and different chromosomes.

In conclusion, our study challenges the conventional single-target paradigm in CLL therapy, advocating for a higher-level, network-oriented strategy. The identification and hierarchical ranking of 20-30 significant proteins, amidst the roughly 20,000 synthesized by the human organism, represent a leap in signal detection and amplification [73]. This nuanced profiling, achieved via a statistical thermodynamics approach, underscores the potential of targeting a selective array of 5-6 network nodes. This selectivity is crucial to mitigate the risk of adverse effects caused by overlapping off-target interactions commonly seen with broader therapeutic targets. The proteins highlighted in our research, notably within the Wnt signaling pathway, are not merely isolated entities but components of a complex network that drives the CLL pathology. Therefore, our proposed method does not end at the identification of these proteins but extends to rank-ordering them in terms of therapeutic relevance. The next step for validating the findings involves experimental assays using siRNA [74] or small molecule inhibitors, which will provide the empirical backbone for our theoretical model. Such an approach may revolutionize the current treatment regimens by transitioning from a one-size-fits-all model to a more customized, patientspecific strategy. This could be especially beneficial given the genetic variability among CLL patients, as indicated by the inconsistent mutation patterns observed in whole-genome sequencing. By incorporating the principles of systems biology and acknowledging the network dynamics of protein interactions, we can begin to envision a more effective, personalized therapeutic landscape for CLL. This, in turn, may pave the way for similar strategies in other cancers, marking a paradigm shift in oncological treatment towards precision medicine.

## Appendix 1 Notes on the genomic location of key genes “involved” in CLL (from GeneCards.org)

### BTK is found in the X chromosome

*Genomic Locations for BTK Gene*

*chrX:101,349,447-101,390,796*

(GRCh38/hg38)

*Size:*

41,350 bases

*Orientation:*

Minus strand

*chrX:100,604,435-100,641,212*

(GRCh37/hg19)

*Size:*

36,778 bases

*Orientation:*

Minus strand

*Genomic View for BTK Gene*

Genes around BTK on UCSC Golden Path with <u>GeneCards custom track</u>

*Cytogenetic band:*

Xq22.1 by <u>HGNC</u>

Xq22.1 by <u>Entrez Gene</u>

Xq22.1 by <u>Ensembl</u>**BTK Gene in genomic location: bands according to Ensembl, locations according to** GeneLoc (and/or Entrez Gene and/or Ensembl if different)

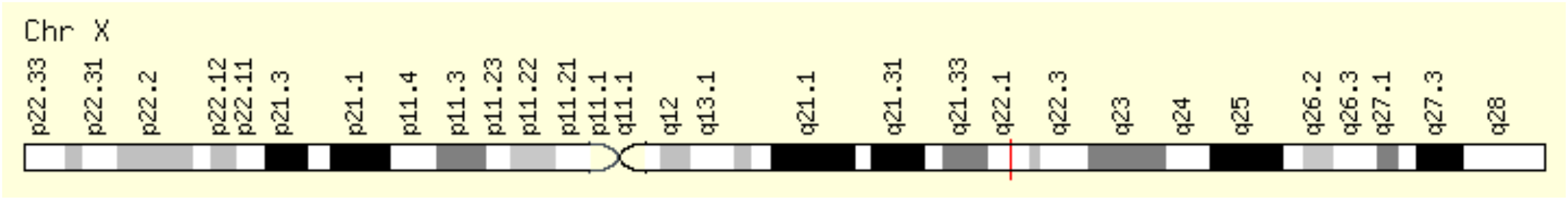

### NFKB1 is in chromosome 4

*Genomic Locations for NFKB1 Gene*

chr4:102,501,329-102,617,302

(GRCh38/hg38) Size:

115,974 bases Orientation:

Plus strand

chr4:103,422,486-103,538,459

(GRCh37/hg19) Size:

115,974 bases Orientation:

Plus strand

Genomic View for NFKB1 Gene

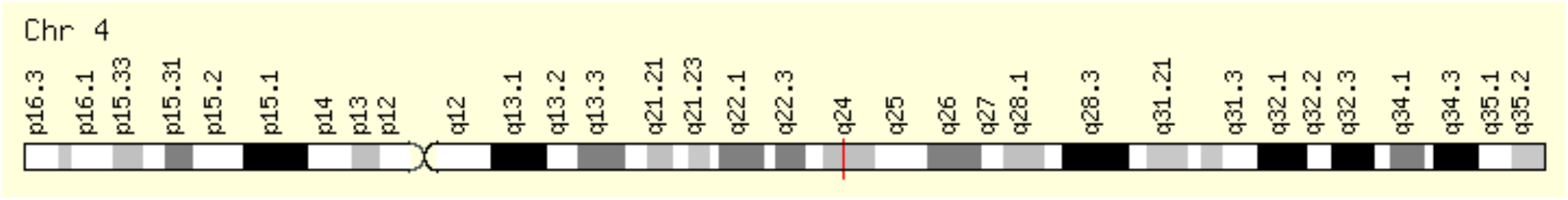

### NFKB2 is on chromosome 10

*Genomic Locations for NFKB2 Gene*

chr10:102,394,110-102,402,529

(GRCh38/hg38) Size:

8,420 bases Orientation:

Plus strand

chr10:104,153,867-104,162,281

(GRCh37/hg19) Size:

8,415 bases Orientation:

Plus strand

Genomic View for NFKB2 Gene

Genes around NFKB2 on UCSC Golden Path with <u>GeneCards custom track</u>

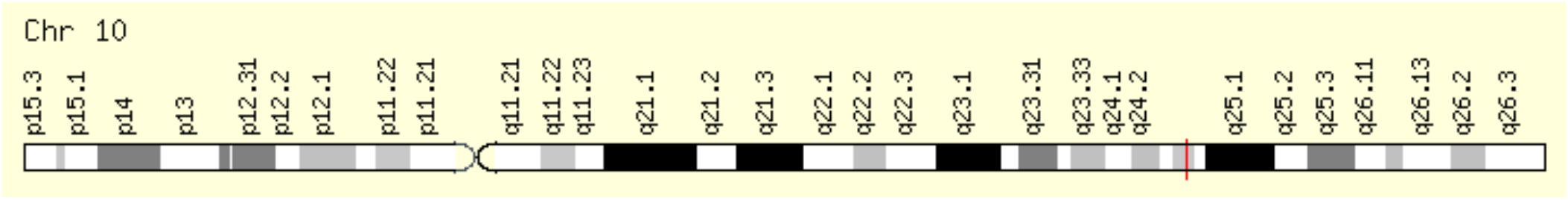

### JAK1 is on chromosome 1

*Genomic Locations for JAK1 Gene*

chr1:64,833,223-65,067,754

(GRCh38/hg38) Size:

234,532 bases Orientation:

Minus strand

chr1:65,298,906-65,432,187

(GRCh37/hg19) Size:

133,282 bases Orientation:

Minus strand

Genomic View for JAK1 Gene

Genes around **JAK**1 on UCSC Golden Path with <u>GeneCards custom track</u>

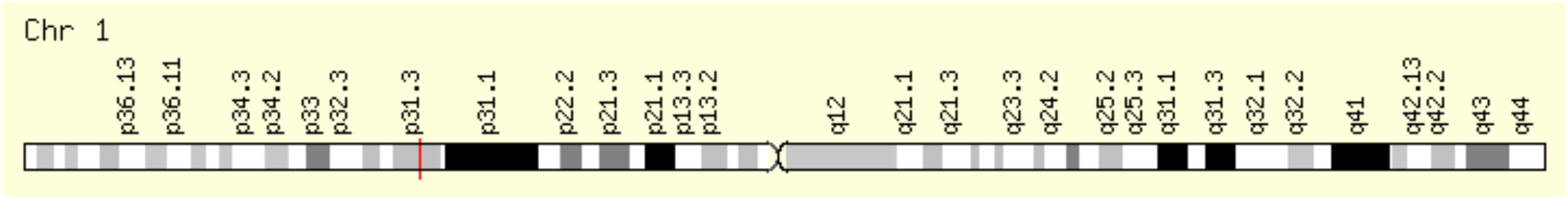

### JAK2 is on chromosome 9

*Genomic Locations for JAK2 Gene*

chr9:4,984,390-5,128,183

(GRCh38/hg38) Size:

143,794 bases Orientation:

Plus strand

chr9:4,985,033-5,128,183

(GRCh37/hg19) Size:

143,151 bases Orientation:

Plus strand

Genomic View for JAK2 Gene

Genes around **JAK**2 on UCSC Golden Path with <u>GeneCards custom track</u>

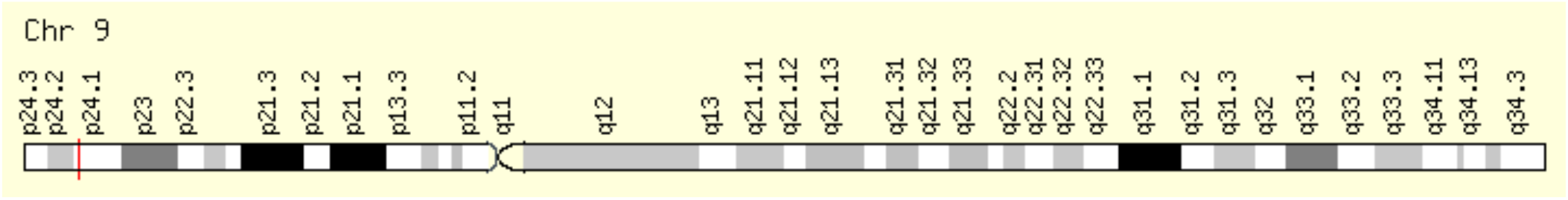

### JAK3 is on chromosome 19

*Genomic Locations for JAK3 Gene*

Genomic Locations for **JAK**3 Gene

chr19:17,824,780-17,848,071

(GRCh38/hg38)

Size:

23,292 bases

Orientation:

Minus strand

chr19:17,935,589-17,958,880

(GRCh37/hg19)

Size:

23,292 bases

Orientation:

Minus strand

Genomic View for JAK3 Gene

Genes around **JAK**3 on UCSC Golden Path with <u>GeneCards custom track</u>

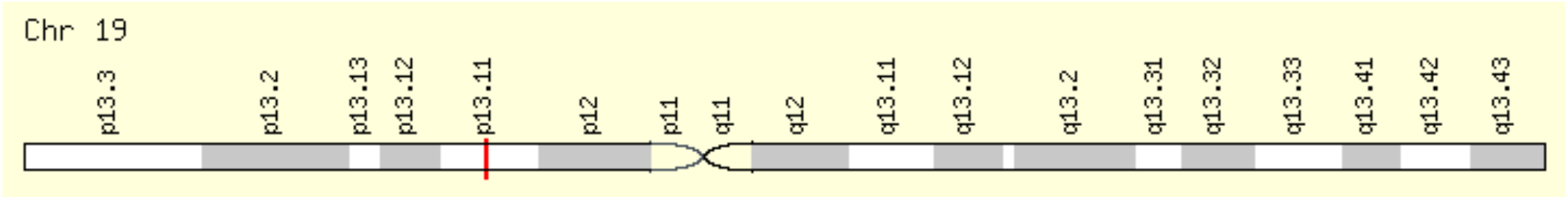

### STAT1 is on chromosome 2

*Genomic Locations for STAT1 Gene*

Genomic Locations for STAT1 Gene

chr2:190,964,358-191,020,960

(GRCh38/hg38)

Size:

56,603 bases

Orientation:

Minus strand

chr2:191,829,084-191,885,686

(GRCh37/hg19)

Size:

56,603 bases

Orientation:

Minus strand

Genomic View for STAT1 Gene

Genes around STAT1 on UCSC Golden Path with <u>GeneCards custom track</u>

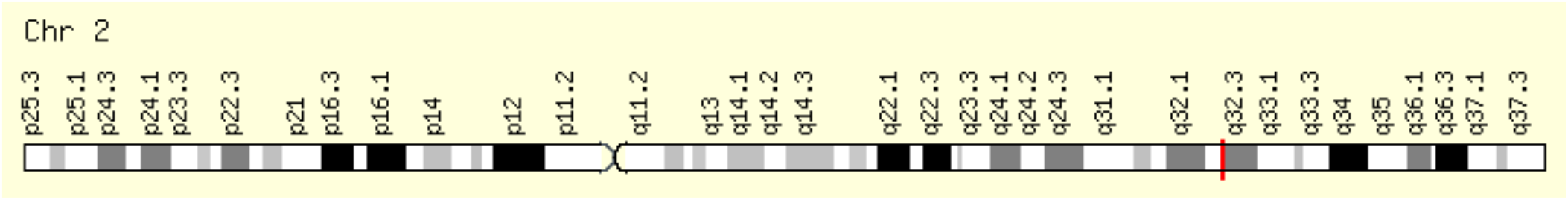

### STAT2 is on chromosome 12

*Genomic Locations for STAT2 Gene*

Genomic Locations for STAT2 Gene

chr12:56,341,597-56,360,253

(GRCh38/hg38)

Size:

18,657 bases

Orientation:

Minus strand

chr12:56,735,381-56,754,037

(GRCh37/hg19)

Size:

18,657 bases

Orientation:

Minus strand

Genomic View for STAT2 Gene

Genes around STAT2 on UCSC Golden Path with <u>GeneCards custom track</u>

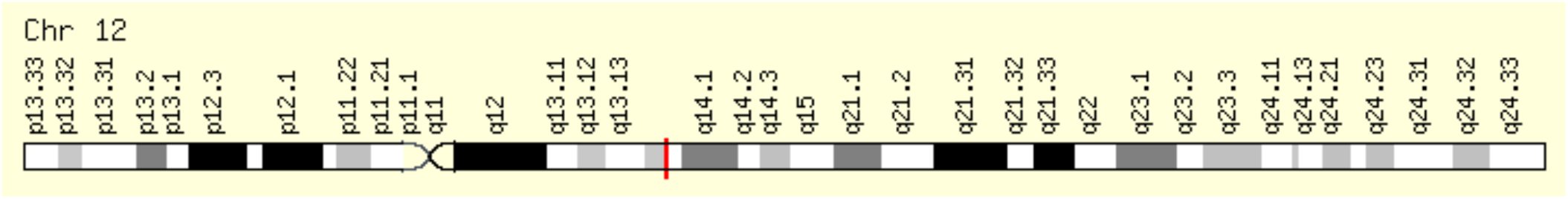

### STAT3 is on chromosome 17

*Genomic Locations for STAT3 Gene*

Genomic Locations for STAT3 Gene

chr17:42,313,324-42,388,568

(GRCh38/hg38)

Size:

75,245 bases

Orientation:

Minus strand

chr17:40,465,342-40,540,586

(GRCh37/hg19)

Size:

75,245 bases

Orientation:

Minus strand

Genomic View for STAT3 Gene

Genes around STAT3 on UCSC Golden Path with <u>GeneCards custom track</u>

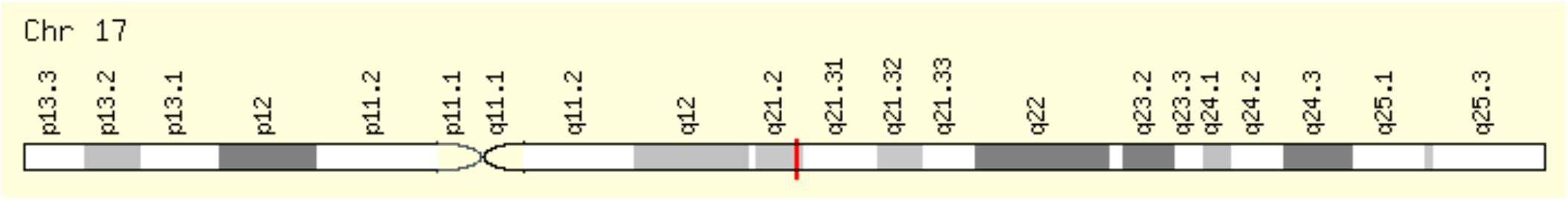

### NOTCH1 is on chromosome 9

*Genomic Locations for NOTCH1 Gene*

Genomic Locations for NOTCH1 Gene

chr9:136,494,433-136,546,048

(GRCh38/hg38)

Size:

51,616 bases

Orientation:

Minus strand

chr9:139,388,896-139,440,314

(GRCh37/hg19)

Size:

51,419 bases

Orientation:

Minus strand

Genomic View for NOTCH1 Gene

Genes around NOTCH1 on UCSC Golden Path with <u>GeneCards custom track</u>

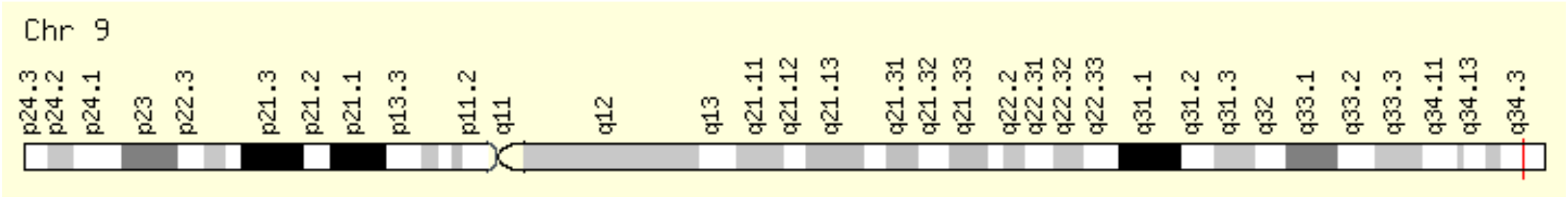

### BCL2 is on chromosome 18

*Genomic Locations for BCL2 Gene*

Genomic Locations for BCL2 Gene

chr18:63,123,346-63,320,128

(GRCh38/hg38)

Size:

196,783 bases

Orientation:

Minus strand

chr18:60,790,579-60,987,361

(GRCh37/hg19)

Size:

196,783 bases

Orientation:

Minus strand

Genomic View for BCL2 Gene

Genes around BCL2 on UCSC Golden Path with <u>GeneCards custom track</u>

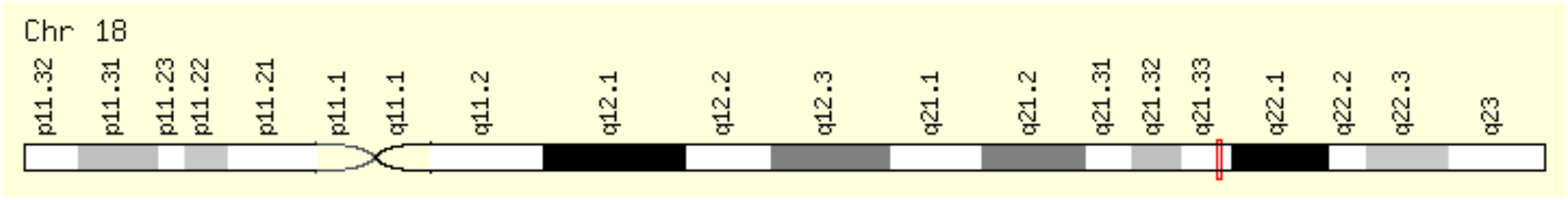

### EEF2 is on chromosome 19

*Genomic Locations for EEF2 Gene*

Genomic Locations for EEF2 Gene

chr19:3,976,056-3,985,463

(GRCh38/hg38)

Size:

9,408 bases

Orientation:

Minus strand

chr19:3,976,054-3,985,467

(GRCh37/hg19)

Size:

9,414 bases

Orientation:

Minus strand

Genomic View for EEF2 Gene

Genes around EEF2 on UCSC Golden Path with <u>GeneCards custom track</u>

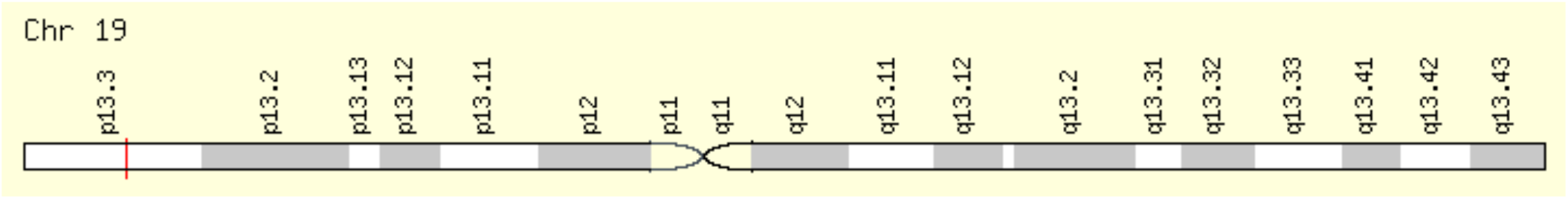

### ABL1 is in chromosome 9

*Genomic Locations for ABL1 Gene*

Genomic Locations for ABL1 Gene

chr9:130,713,016-130,887,675

(GRCh38/hg38)

Size:

174,660 bases

Orientation:

Plus strand

chr9:133,589,268-133,763,062

(GRCh37/hg19)

Size:

173,795 bases

Orientation:

Plus strand

Genomic View for ABL1 Gene

Genes around ABL1 on UCSC Golden Path with <u>GeneCards custom track</u>

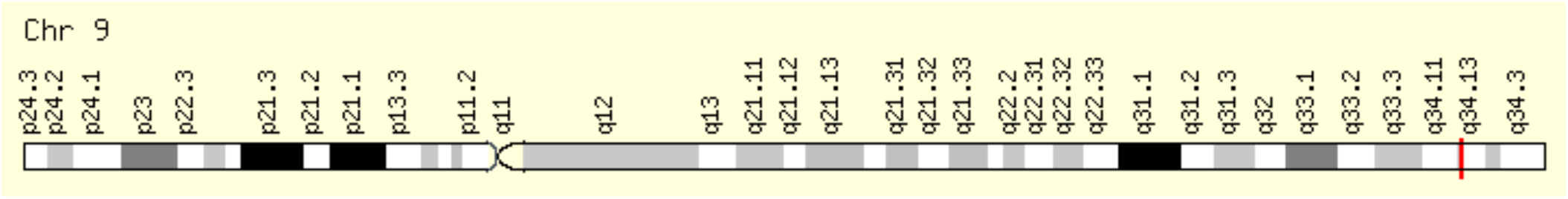

### BCR3 (BCRP3) is in chromosome 22

*Genomic Locations for BCRP3 Gene*

Genomic Locations for BCRP3 Gene

chr22:24,632,915-24,653,360

(GRCh38/hg38)

Size:

20,446 bases

Orientation:

Plus strand

chr22:25,028,882-25,049,327

(GRCh37/hg19)

Size:

20,446 bases

Orientation:

Plus strand

Genomic View for BCRP3 Gene

Genes around BCRP3 on UCSC Golden Path with <u>GeneCards custom track</u>

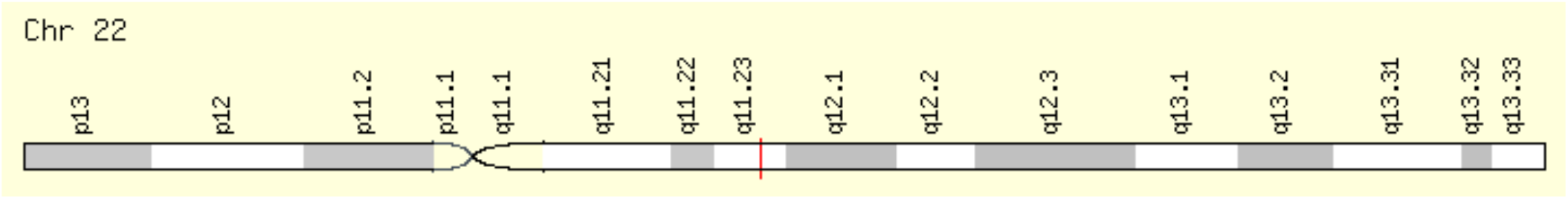

### SF3B1 is in chromosome 2

*Genomic Locations for SF3B1 Gene*

Genomic Locations for SF3B1 Gene

chr2:197,388,515-197,435,091

(GRCh38/hg38)

Size:

46,577 bases

Orientation:

Minus strand

chr2:198,254,508-198,299,815

(GRCh37/hg19)

Size:

45,308 bases

Orientation:

Minus strand

Genomic View for SF3B1 Gene

Genes around SF3B1 on UCSC Golden Path with <u>GeneCards custom track</u>

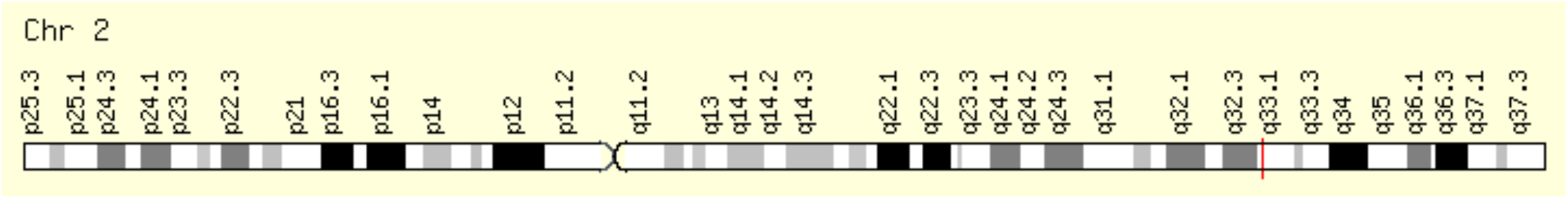

### ZAP70 is on chromosome 2

*Genomic Locations for ZAP70 Gene*

Genomic Locations for ZAP70 Gene

chr2:97,713,560-97,744,327

(GRCh38/hg38)

Size:

30,768 bases

Orientation:

Plus strand

chr2:98,330,023-98,356,325

(GRCh37/hg19)

Size:

26,303 bases

Orientation:

Plus strand

Genomic View for ZAP70 Gene

Genes around ZAP70 on UCSC Golden Path with <u>GeneCards custom track</u>

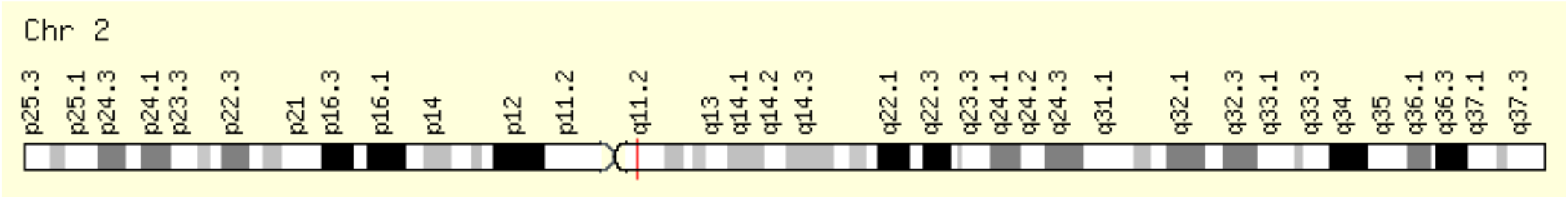

### MYC is on chromosome 8

*Genomic Locations for MYC Gene*

Genomic Locations for MYC Gene

chr8:127,735,434-127,742,951

(GRCh38/hg38)

Size:

7,518 bases

Orientation:

Plus strand

chr8:128,747,680-128,753,680

(GRCh37/hg19)

Size:

6,001 bases

Orientation:

Plus strand

Genomic View for MYC Gene

Genes around MYC on UCSC Golden Path with <u>GeneCards custom track</u>

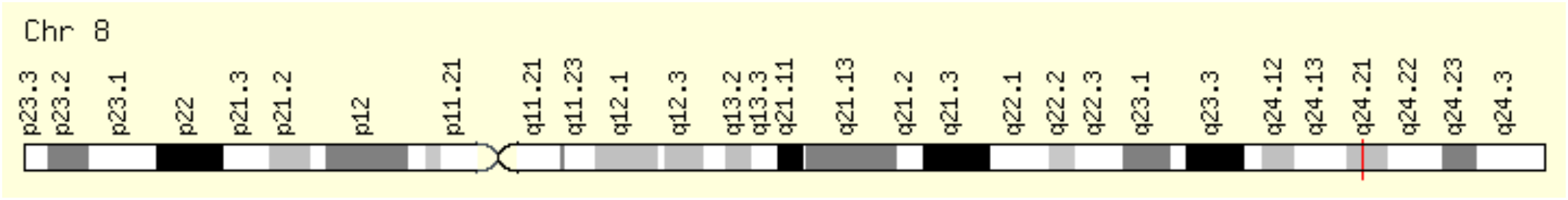

## Appendix 2. Union of the Published Gene List Reviewed in this Study

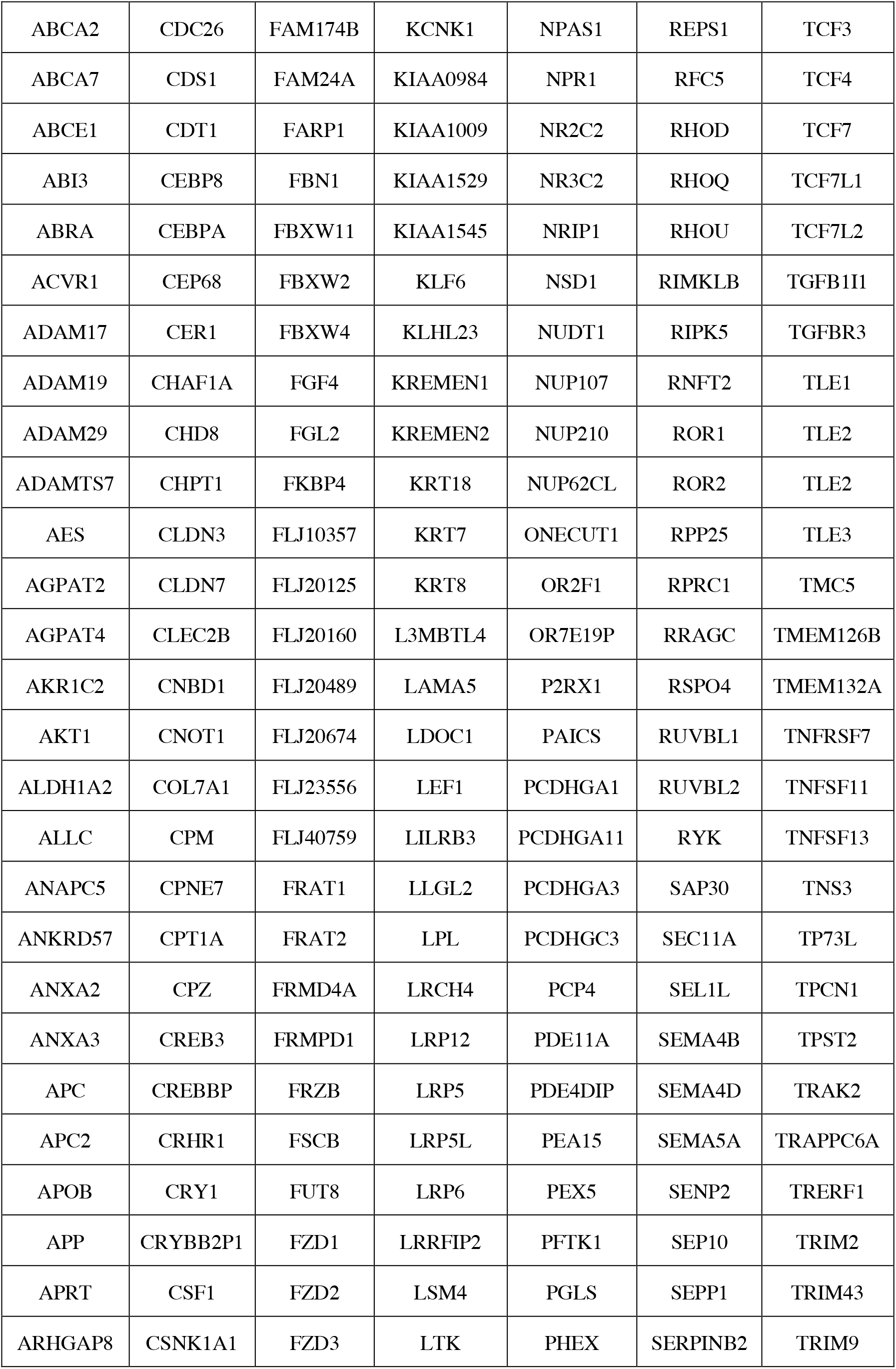
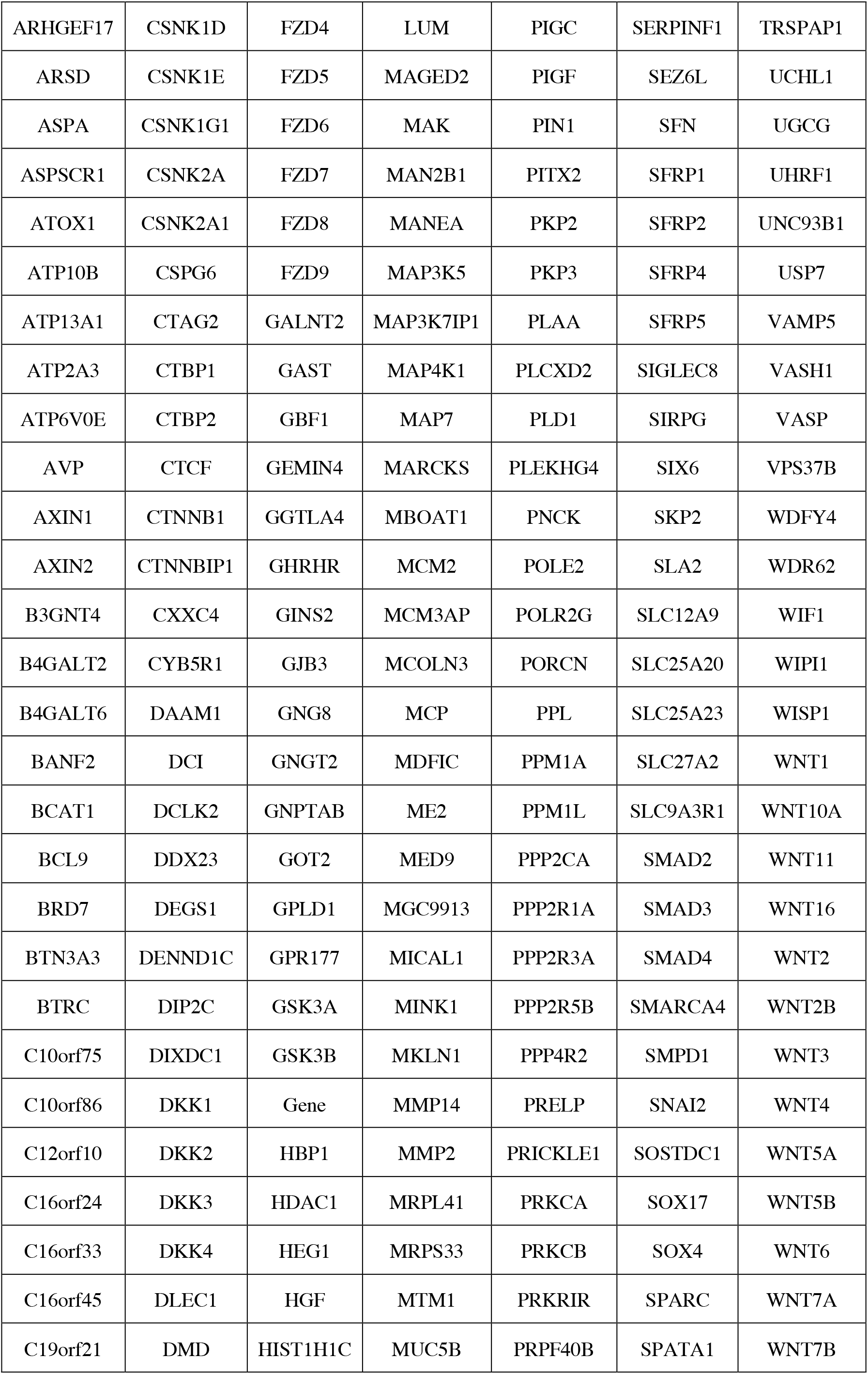
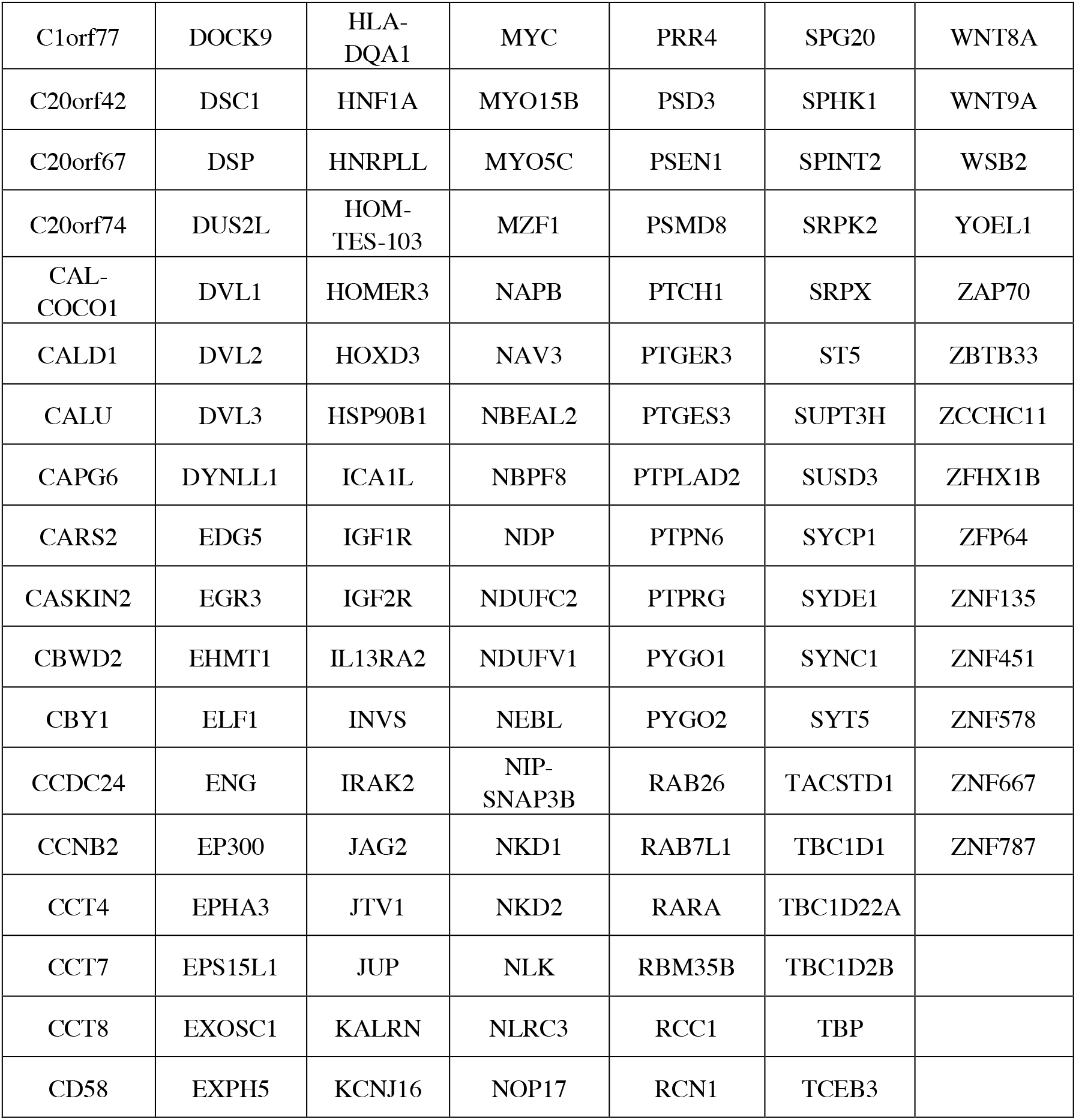

## Appendix 3. Integrative Analysis of CLL by t-SNE Visualization

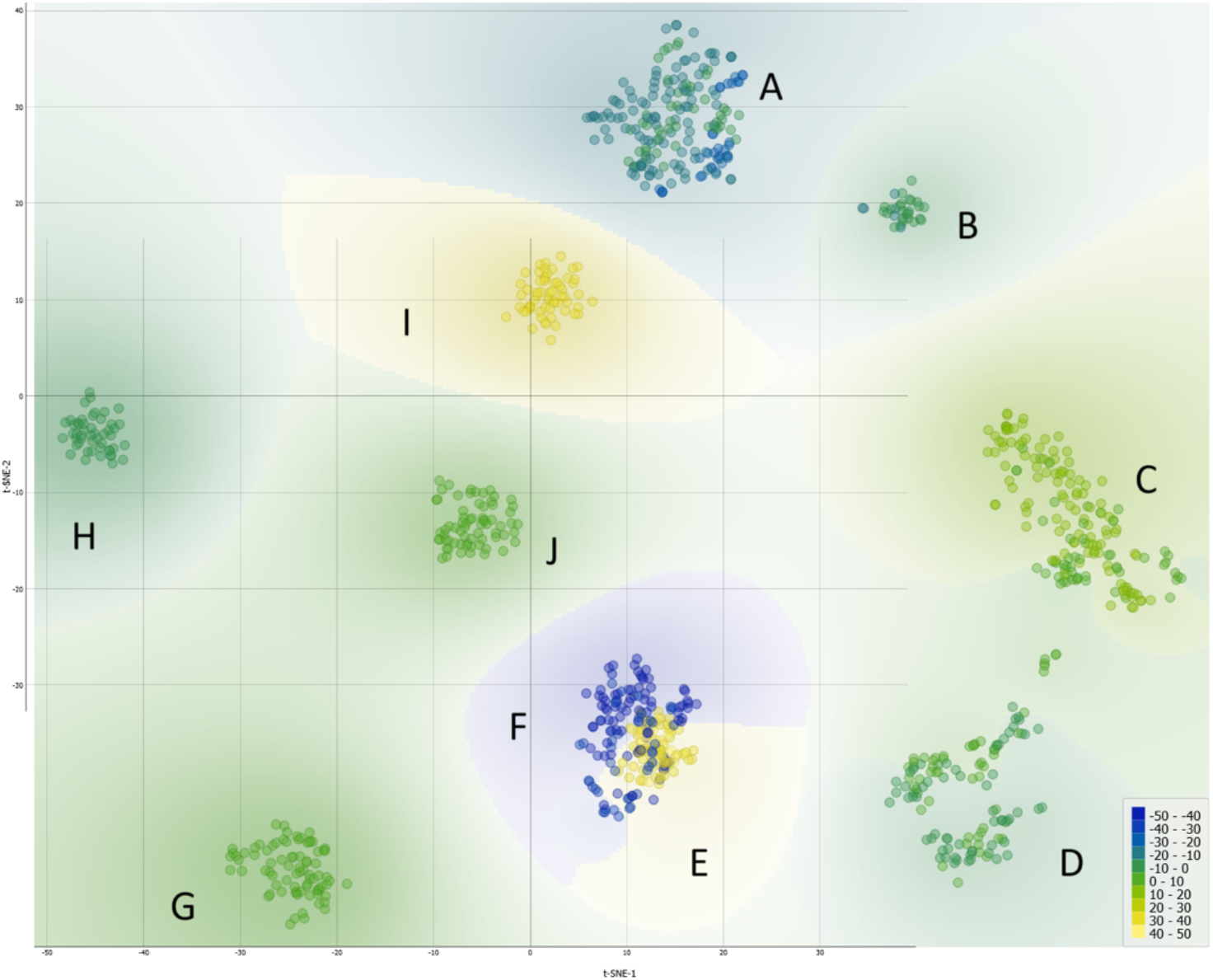

**t-Distributed Stochastic Neighbor Embedding (t-SNE) Analysis of CLL Samples and Subgroups**. The visualization displays a non-linear dimensionality reduction of the complex gene expression data, with each point representing individual samples. The layout highlights the nuanced relationships and 10 clusters labeled from A to J within the CLL dataset consisted of the 1001 patients, uncovering uncaptured subtleties through the network analysis. Samples included in the dataset are either diagnosed CLL patients (sick) or wild-type patients (normal) without CLL.

- Cluster A encapsulates a significant cohort focusing on the Wnt signaling pathway, a key player in CLL pathogenesis, with a total of 179 patients. Within this cluster, 21 are wild type, while 158 are CLL patient samples, all derived from the GSE31048 dataset. This dataset offers a unique look at both normal and CLL-affected B cells, allowing for a direct comparison of Wnt pathway gene expression and Wnt-regulated gene expression. The marked difference in numbers between the normal (12 and 9, respectively) and sick (149 and 9, respectively) groups underscores the aberrant expression within CLL-affected B cells, highlighting the pathway’s prominence in these cellular states. The study’s in-depth focus on the Wnt pathway is well-founded, as it is pivotal in cellular processes that are often disrupted in CLL, thus potentially illuminating new therapeutic avenues.
- Cluster B includes a smaller, yet focused subset of 42 patients, of which 12 are wild type and 30 are CLL patients. This cluster continues the examination of the Wnt pathway’s role in CLL as part of the GSE31048 study, indicative of a unique or divergent role of Wnt signaling in this subset. It is particularly noteworthy that this dataset includes expression data from CLL B cells with and without Wnt3a treatment, providing insights into the pathway’s functionality and potential for targeted therapies. The comparative analysis of Wnt gene expression between the normal and CLL B cells offers additional evidence for the pathway’s critical role in the disease process.
- Cluster C, with 136 patients, 24 normal and 112 sick, merges data from two distinct studies, GSE10139 and GSE50006, providing a broader scope by incorporating a genomic approach from GSE10139 to improve prognosis and therapeutic response predictions, and juxtaposes this with expression data from CLL tumors and healthy donor B cells from GSE50006. The inclusion of CLL-blood samples enriches the dataset, illustrating the heterogeneity within CLL and potentially reflecting different disease phases or subtypes. The blend of these datasets furnishes a more comprehensive understanding of the disease, highlighting the heterogeneity of CLL and reinforcing the necessity of personalized medicine approaches.
- Cluster D presents a cohort of 152 patients, predominantly sick (144) with a small representation of normal B cells (8), combining data from GSE10139 and GSE50006. This distribution, primarily composed of CLL and CLL-blood samples, continues to emphasize the genetic and expression-level diversity found in CLL, supporting the need for in-depth analysis to discern the nuances of the disease’s progression and the potential response to treatments.
- Cluster E is a homogeneous group consisting entirely of 100 sick patients from the GSE49896 dataset. This study spotlights the microRNA-150’s influence on B-Cell Receptor signaling by modulating GAB1 and FOXP1 gene expressions, which are implicated in CLL. MicroRNAs are crucial post-transcriptional regulators, and their role in CLL adds an additional layer to understanding the disease’s complexity and potential intervention points.
- In Cluster F, 130 CLL patients from the GSE39671 dataset are studied, all of whom have undergone treatment. The data represent a temporal progression, with sampling times to first treatment recorded, allowing for an exploration of the disease’s evolution over time. The dataset’s analysis provides prognostic subnetworks which can help predict disease progression and highlight the converging pathways in CLL, opening new avenues for tailored treatments.
- Cluster G, comprising 75 CLL patients from the GSE69034 study, delves into the gene expression profiles linked with the MYD88 L265P mutations in conjunction with IGHV mutation status. The presence of the MYD88 L265P mutation, a notable variant found within the MYD88 gene that encodes a key adaptor protein in the Toll-like receptor and IL-1 receptor pathways, has been tied to specific prognostic outcomes in CLL. This mutation is known to activate downstream signaling pathways aberrantly, which can contribute to the uncontrolled proliferation of B cells characteristic of CLL. The dataset’s inclusion in the study facilitates a detailed investigation into the mutation’s role and its pathway associations in CLL, offering a potential explanation for the varying responses to treatment observed in patient populations. By analyzing the gene expression patterns influenced by the MYD88 L265P mutation alongside the IGHV mutation status, a well-established prognostic marker in CLL, it unravels the complex interplay between genetic aberrations and their impact on the disease’s clinical course. The correlation between MYD88 L265P mutations and factors such as treatment resistance, disease progression, and overall survival can be assessed. This is particularly crucial, as the mutation’s impact on signaling pathways may suggest new therapeutic targets or strategies for intervention. Groundbreaking biomarkers are likely to be identified for early detection and prognosis by understanding the biological context in which these mutations operate, while also highlighting the therapeutic relevance of targeting the MYD88 pathway in certain subsets of CLL patients, which implies the importance of precision medicine in the management of CLL. Based on the insights into the specific mutations driving the disease in individual patients, therapies can be customized to target these genetic abnormalities more effectively. In the case of MYD88 L265P, its presence could signify a need for targeted inhibitors that can mitigate its downstream effects, thereby introducing a new dimension to personalized CLL treatment paradigms.
- Cluster H is a cohort of 84 CLL patients from GSE28654, all carrying the IgVHMT mutation and exhibiting negative ZAP-70 expression. The absence of ZAP-70 expression, a kinase linked to CLL, together with the mutational profile, provides a critical connection for investigation. This relationship implicates the substantial impact of the mutation on CLL’s clinical progression and pinpoints the need for detailed genetic analysis in crafting specialized treatments.
- In Cluster I, 28 sick patients from GSE28654 are categorized by the presence of the IgVHUM mutation and positive ZAP-70 expression, helping to understand the disease’s heterogeneity since ZAP-70 positivity is often linked with a more aggressive CLL form. The combination of mutational status and ZAP-70 expression levels provides valuable prognostic information.
- The expression of ZAP-70 in CLL and its relevance as a molecular marker is particularly illuminating. For Cluster H, the collective profile of CLL patients characterized by the IgVHMT mutation yet displaying an absence of ZAP-70 expression represents a subset where traditional prognostic markers may predict a more favorable clinical course. In the broader landscape of our findings, this cluster could suggest that ZAP-70’s negativity may reflect a less aggressive form of CLL, where the malignant B cells might not engage in the same signaling pathways that are characteristic of more virulent variants. Consequently, these insights bolster the argument for personalized therapeutic approaches, enabling clinicians to tailor treatments to the specific molecular makeup presented by individual CLL cases. Conversely, patients in Cluster I, characterized by the IgVHUM mutation concomitant with positive ZAP-70 expression, suggest a more aggressive manifestation of the disease. This association aligns with the understanding that ZAP-70 positivity mirrors the behavior of unmutated IgVH status, commonly linked to a robust disease progression and a less favorable response to conventional therapies. Here, ZAP-70 serves not just as a prognostic marker but potentially as a therapeutic target, whereby modulation of its expression or function could impact CLL cell survival. This reiterates the substantial role that ZAP-70 plays in CLL. It acts as a bifurcation point in the disease’s prognostic roadmap, where its expression could either denote a need for more aggressive treatment or suggest a less intensive therapeutic course. The interplay of ZAP-70 with IgVH mutation status, as demonstrated in our clusters, provides a clearer understanding of disease heterogeneity and patient stratification. The overall results of the study thus advocate for the integration of ZAP-70 status into prognostic models and therapeutic decision-making algorithms, emphasizing its contribution not only to prognostication but potentially to the development of targeted CLL therapies.
- Cluster J, mirroring Cluster G, includes another set of 75 CLL patients from the GSE69034 dataset, indicating the significant role of MYD88 L265P mutations in CLL, providing a robust dataset for the exploration of mutation-associated gene expression patterns and their prognostic significance.

The heterogeneity of gene mutations across CLL patients underscores the intricate complexity of this malignancy, accentuating the necessity for individualized therapeutic strategies. The disparities unearthed by t-SNE analysis manifest in the distinct molecular signatures differentiating normal B cells from CLL-B cells, which reflect divergent evolutionary trajectories within the disease’s progression. Notably, the aberrant expression of Wnt pathway genes in CLL cells, as revealed by our cluster analysis, pinpoints this pathway’s pivotal role in CLL pathobiology. The presence of specific gene expressions within clusters, particularly those highlighted by the t-SNE method (clusters A and B), points to the pathway’s disrupted regulation, which is suggestive of patient-specific disease mechanisms that contribute to CLL’s heterogeneity. Simultaneously, the funding emphasizes that alterations in the Wnt signaling pathway are not universally present but vary among patients, reinforcing the pathway’s contribution to the disease complexity. The therapeutic potential of targeting Wnt pathway proteins is corroborated by their identified roles in vital cellular functions, with their significance accentuated by Betti number estimates, which propose these proteins as central players in CLL’s pathogenesis rather than inconsequential elements. Such insights solidify the imperative for a more comprehensive, multi-scalar study from cellular to genomic dimensions to forge ahead with personalized treatments for CLL.

## Author Contributions

GLK suggested the study. EAR and GLK selected the datasets. EAR conducted the computational study and wrote the first draft with input from GLK. HTS supported EAR and provided consulting on the project. GP contributed the study on drugs and rewrote the paper with the assistance of JZ. HH performed the clustering analysis. JAT contributed to the cancer drug analysis and wrote the paper with GP and EAR.

## Data Availability Statement

The data for the CLL study is from the GEO Website and more specifically are the following datasets: GSE10139, GSE28654, GSE31048, GSE39671, GSE49896, GSE50006, and GSE69034.

## Acknowledgments

JAT acknowledges the funding support received from NSERC (Canada) for this project.

